# Potential of Local *Bacillus* spp. Isolates as Wilt Disease Biocontrol Agents for *Fusarium* (*Fusarium oxysporum* f. sp. *cepae*) on Wakegi Onions (*Allium* × *wakegi* Araki)

**DOI:** 10.1101/2023.03.28.534608

**Authors:** Asrul Asrul

## Abstract

The use of biological agents as a biocontrol against plant pathogens was often ineffective because it worked slowly. The objective of this research was to examine the potential of local isolates of Bacillus spp. as a biocontrol agent in suppressing Fusarium wilt disease (*Fusarium oxysporum* f. sp. *cepae*) on wakegi onions. The research was designed using a completely randomized design with the treatment of rhizosphere bacterial isolates. The treatments consisted of control (without isolate application), isolates KP17, KP5, DB9, DB12, DB18, DG4, and DG11 so that the number of treatments was eight. Each treatment was repeated 5 times and each replication consisted of 10 wakegi onion plants. This research was divided into 2 stages, namely laboratory research which included isolation, characterization of colony morphology of rhizosphere bacterial isolates, and in vitro testing of the inhibitory power of biocontrol agents against pathogens. The test in the greenhouse was in the form of a disease case suppression test. The results obtained seven candidate isolates of biocontrol from 46 isolates obtained from the rhizosphere of the wakegi onion. These isolates had similarities with Bacillus spp. based on colony morphology, physiology, and biochemistry characteristics. Among the isolates found, the DB12 isolate had the potential to be developed as a biocontrol agent compared to other isolates.

## Introduction

Wakegi onion (*Allium* × *wakegi* Araki) is one of the leading vegetable commodities in Central Sulawesi, and it has been widely cultivated and developed by farmers, especially in the Sigil Regency. The people of Central Sulawesi know wakegi onions as fried onions, while shallots were mostly used as cooking spices. Shallots have the characteristic of being able to flower in the lowlands or highlands, on the contrary, wakegi onions cannot flower. Wakegi onion is the result of a natural interspecific cross between leek (*Allium fistulosum* L.) and shallot (*Allium cepa* L. *aggregatum* group) [1]. The wakegi onion cultivars include ‘Palasa’, ‘Tinombo’, ‘Palu Valley’ (Central Sulawesi), and ‘Sumenep’ (Madura, East Java) [2].

One of the main obstacles in the development of wakegi onion cultivation in Sigi Regency was the presence of *Fusarium* wilt disease/basal rot of stems or known as moler disease caused by the fungus *Fusarium oxysporum* Schlechtend [3]. Foc (*Fusarium oxysporum* Schlechtend) fungus attacks can cause yield losses of up to 50% [4] and are even capable of causing crop failure if the environment was supportive of fungi [5]. This situation was often complained of by local farmers; especially when entering the rainy season, the risk of pathogen attack was higher so it had to be handled immediately. Therefore, the presence of the disease could not be ignored, because even though the Foc attack was rather slow, if onions were infected it could cause plant death.

Foc fungus control by farmers generally still involves synthetic fungicides whose use could cause various negative effects, especially pesticide residues that are harmful to the environment and human health [6,7]. Residues left in agricultural products often worry people who consume foods such as vegetables and fruits because they could endanger health. The use of Foc-resistant varieties was generally more practical, effective, and environmentally friendly, but there were not many available varieties that had resistance to these pathogens.

The use of biological agents is currently getting attention from farmers because it is economically cheaper and more effective, safer for farmers and the environment, and maintains ecological balance. However, the biological agents generally applied were specific, so they could not last long and were difficult to develop in an environment that was not their natural habitat. Therefore, it was necessary to develop control technology that was adaptive to the local ecosystem and sustainable based on the principle of “from nature for nature” by utilizing antagonistic local isolates of rhizosphere bacteria (indigenous). This was because local bacterial isolates were more adaptive to survive and had better potential to suppress pathogens in their origin area compared to using isolates introduced from other areas [8]. According to Erwanti et al. [9], the use of local antagonist microbes to control pathogens in the area would provide better and more effective results than antagonistic microbes introduced from outside.

One of the potential rhizosphere bacteria that could be recommended as a substitute for synthetic pesticides and was widely used as a biological control agent was *Bacillus* spp. This was because these bacteria could suppress the development of several important pathogenic fungi such as *Fusarium oxysporum, Phytophthora, Pythium, Alternaria, Colletotrichum, Plasmodiophora brassicae* [10], *Phytophthora infestans* [11], *F. graminearum, F. subglutinans,* and *F. verticilliodes, F. solani* [12], and *Rhizoctonia solani* [13].

The suppression by *Bacillus* spp. of the growth of pathogenic fungi was done through antagonistic mechanisms such as antibiosis (antibiotics), enzyme production, competition (space and nutrition), parasitism (bacteria moved to germinated spores and adhered to the surface through polarity, then secreted catalytic enzymes), induced systemic resistance in plants (through the jasmonic acid pathway) and siderophore production [14,15].

Among these antagonistic mechanisms, antibiosis was the main mechanism for *Bacillus* spp. bacteria to inhibit the growth of pathogens through the production of antifungal compounds (antibiotics) [12] that were toxic to microbes. The antibiotics produced include bacillomycin, mycobacillin, fungistatin, iturin, fengisin, plipastatin, surfactin, bacilisin, lipopeptide, biosurfactant, penazine, and hydrogen cyanide (HCN) [15]. *Bacillus* spp. bacteria also produced enzymes that could degrade (break down) the cell walls of pathogenic fungi so that their growth was inhibited and the fungus dies, including chitinase, protease, b-1,3 glucanase, and cellulase, and stimulated the body’s defenses through induction of systemic resistance through the jasmonic acid pathway, salicylic acid [14,16]. In addition, it also produced siderophores such as pioselin and pioverdin to inhibit the proliferation of phytopathogens by chelating iron (Fe3+) in the rhizosphere area so that it became unavailable to pathogens [15]. The ability to produce antibiotics, enzymes, siderophores, and the induction of resistance made *Bacillus* spp. bacteria able to act as biopesticides. Meanwhile, the ability of antagonistic bacteria to fix nitrogen, dissolve phosphate to be absorbed by plants, and produce growth hormone was the role of these bacteria as biofertilizers.

So far, knowledge about morphological characters, inhibition mechanisms, and effectiveness of local isolates of antagonistic bacteria that would be used to control Foc fungi was unknown. Therefore, it was necessary to carry out isolation and characterization activities to find specific and adaptive biocontrol agents to apply environmentally friendly disease control techniques. Furthermore, isolates could be developed an effective control method against Foc pathogens by using a *Bacillus* spp. biocontrol agent. This research aimed to examine the potential of local isolates of *Bacillus* spp. as a biocontrol agent in suppressing *Fusarium* wilt disease in wakegi onions.

## Materials and methods

This research was done from April to September 2021. The research was divided into two stages of activity. The first stage was in the laboratory: isolation, hypersensitivity reaction test, characterization of rhizosphere bacterial isolates based on colony morphology, physiology, and biochemistry, and inhibition power test. The second stage was in the greenhouse and included suppression tests of bacterial isolates against *Fusarium* wilt disease in wakegi onion plants.

### Research Preparation

The tools and materials used during the research included: 0.9 mm hole sieve, microscope, LAF (Laminar Air Flow), autoclave, hand spray, hemocytometer, micropipette, test tube, Erlenmeyer flask, glass spatula, watch-glass, Petri dish, fungi *F. oxysporum* f. sp. *cepae*, healthy wakegi onion bulbs, sterile water, toothpicks, Potato Dextrose Agar (PDA) and Sodium Agar (NA) media, and 75% alcohol.

### Rhizosphere Soil Sampling

Soil samples were obtained from the rhizosphere layer of healthy wakegi onions located in Petobo Village (Palu City), Bulupontu/Sidera Village (Sigi Regency), and Guntarano Village (Donggala Regency). Soil samples were taken using a small shovel in the area around the plant roots. Next, the sample was wrapped in wet paper and put in a plastic bag, and stored in an ice flask to be brought to the laboratory.

### In the Laboratory

#### Isolation of Bacillus spp. from the rhizosphere of the wakegi onion plant

Isolation of antagonistic bacteria was done using the serial dilution method and was specific for *Bacillus* spp. according to [17,18,19]. Cultures were incubated at room temperature for 24 to 48 hours. The grown colonies were then purified using the same medium and tested for hypersensitivity reactions, colony morphology characterization, as well as physiological and biochemical properties testing.

#### Hypersensitivity reaction test

A hypersensitivity reaction (HR) test of antagonistic bacteria isolates to tobacco plants was done according to Asrul et al. [20]. The isolates that caused a negative reaction (no symptoms of necrosis but the leaves remained green) in the inoculated tobacco leaf area were then tested for antagonists to determine their inhibitory power.

#### Characterization of Rhizosphere Bacterial Isolates

Characterization of rhizosphere bacterial isolates was generally done based on the method proposed by Schaad et al. [21]. This characterization aimed to determine the genera of bacterial isolates isolated from the rhizosphere of the wakegi onion. Observation of the morphological characteristics of bacterial colonies of *Bacillus* spp. was done visually on the seventh day after being rejuvenated on NA media. The characterizations included colony morphology (shape, color, margin, elevation, texture, and optical properties), gram reaction, fluorescent pigment, facultative anaerobes, oxidation, catalase, and growth at the temperature of 65°C. The observations were then compared with the morphological characteristics of the colonies previously described by Schaad et al. [21], Calvo & Zuniga [22], Lu et al. [23], Nurcahyanti & Ayu [24], and Prihatiningsih et al. [25].

#### In vitro Inhibition Power Test of Local Isolates

The inhibition power test of rhizosphere bacteria against Foc pathogenic fungi (laboratory collection) was done using dual culture technique according to Jimtha et al. [26]. The inhibitory activity was observed based on the presence of a clear zone formed around the bacterial colonies. Cultures were incubated at room temperature and antagonism interactions were observed at 7 days after inoculation (dai). Percentage of rhizosphere bacteria inhibition was calculated by the formula [27]: P = (R – r)/R × 100%, where P is the percentage of antagonist inhibition power against pathogens, R is the maximum radius of the pathogenic colony stay away from the rhizosphere bacterial colony (cm), and r is the radius of the pathogenic colony opposite with the rhizosphere bacterial colony (cm).

Isolate candidates of rhizosphere bacteria that showed inhibition power greater than 60% were antagonist isolates classified as having effective inhibition activity.

### In the Greenhouse

#### Preparation of Antagonist Bacterial Suspension

The suspension was done aseptically utilizing colonies of rhizosphere bacteria on rejuvenation media aged 48 hours taken using an Ose needle and suspended in a tube containing 10 ml of sterile water. Furthermore, the suspension was made into the mother liquor to calculate the density of the colony population, which was done by the spread dish method. A series of culture tubes containing 9 ml of sterile distilled water were taken, and serial dilutions of 10^-5^, 10^-6^, and 10^-7^ were made. A total of 1 ml from each dilution series was pipetted and grown on NA medium to which 100-ppm cycloheximide was added to suppress fungal growth. Each dilution series was repeated three times. Bacterial colonies that grew were also counted to determine their population density. Next, the suspension was diluted again to obtain a population density of 108 CFU/ml and the bacterial suspension was ready to be applied.

#### Preparation of Pathogenic Fungal Suspension

Preparation of conidia suspension of *F. oxysporum* f. sp. *cepae* was done by taking 1 Ose sample from the mushroom colony that had been rejuvenated, and then putting it in 9 ml of sterile distilled water and homogenizing using a vortex mixer to form a suspension. The formed suspension was calculated for conidia density using a hemocytometer observed under a microscope at a magnification of 400 times. Next, the suspension was diluted to obtain a fungal conidia density of 108 conidia/ml and the suspension was ready to be applied.

#### Fusarium Wilt Suppression Test

Testing of rhizosphere bacterial suppression as a biocontrol agent against *Fusarium* wilt disease on wakegi onions was done in a greenhouse. The research was designed using a completely randomized design (CRD) with the treatment of rhizosphere bacterial isolates. The treatments consisted of control (without isolate application) and isolates KP17, KP5, DB9, DB12, DB18, DG4, and DG11, so that there were eight treatments. Each treatment was repeated five times and each replication consisted of ten wakegi onion plants.

### Treatment Application

The planting medium used is a mixture of soil and manure as basic fertilizer (2:1), which had been sifted and sterilized. The soil was sterilized by tyndallization in a drum with hot steam (temperature of 65°C) for three consecutive days. After the sterilization period was complete, the soil was allowed to cool, then put into a 3-kg polybag. Planting healthy wakegi onions was done by immersing seeds (bulbs) into the soil and carried out in the afternoon.

The disease suppression test was done by growing wakegi onions until they reached the age of 28 days, and then pathogens were applied. Suspension of the pathogen *F. oxysporum* f. sp. *cepae* as much as 10 ml/bulb (density 10^8^ conidia/ml) was poured into planting holes in polybags. A suspension of rhizosphere bacterial isolates, as much as 20 ml/polybag (10^8^ CFU/ml), was introduced into the soil in polybags one week before the inoculation of *F. oxysporum* f. sp. *cepae*. Furthermore, the infestation of the isolate suspension was repeated every 7 days, with up to four total applications. The control treatment was used as a comparison, namely plants without suspension of bacterial isolates.

### Observation

Observation of disease suppression was done on the fifty-sixth day after the infestation of biocontrol agents. The components of the observed pathosystem include the incubation period and the percentage of disease incidence. The incubation period was observed from the first time the pathogen was inoculated until the initial symptoms of fusarium wilt disease appeared. The intensity of the disease was calculated from the onset of symptoms at an interval of 7 days. The percentage of disease incidence was calculated using the formula proposed by Shamyuktha et al. [28] and Lilai et al. [29].

### Data analysis

The data obtained were then analyzed statistically by ANOVA (Analysis of Variance). If there was a significant difference, then the Tukey’s HSD (honestly significant difference) test was performed at the 5% level.

## Results and discussion

### In the Laboratory

A total of 46 bacterial isolates were isolated from the rhizosphere layer of the wakegi onion plant, and after going through a hypersensitivity reaction test, seven local isolates were selected which produced negative reactions. Negative reactions were seen in tobacco leaves injected with local isolate bacterial suspension, which did not cause symptoms of water soaking and necrosis but remained green. The seven local isolates consisted of two isolates (KP17 and KP5) from Petobo Village (Palu City), 3 isolates (DB9, DB12, and DB18) from Bulupontu/Sidera Village (Sigi Regency), and two isolates (DG4 and DG11) from Guntarano Village (Donggala Regency). This negative reaction proved that the local isolate bacteria obtained from the rhizosphere of the wakegi onion were not plant pathogens, but may come from a group of antagonistic bacteria. Mougou & Boughalleb-M’hamdi [30] reported that one of the antagonistic bacteria that reacted negatively to the hypersensitivity test was *Bacillus* spp.

Bacteria that were pathogenic would react positively (compatible) in the hypersensitivity test, which was characterized by the formation of symptoms of necrotic spots on the leaf area where the bacterial suspension was injected. This necrosis symptom was one of the characteristics of a hypersensitive reaction [31]. On the other hand, non-pathogenic bacteria will react negatively (incompatible) by not changing the color of the leaves (remaining green) because the symptoms of necrosis would not appear. Hypersensitivity reactions were a mechanism of plant resistance in response to pathogenic infections in the form of the rapid death of plant tissue cells to localize pathogens so that plants could overcome infections that had the potential to cause disease [21].

The selected local isolates were then tested for in vitro antagonism to determine their inhibition power ability against the growth of the fungal pathogen *F. oxysporum* f. sp. *cepae* (Foc).

### Local Isolate Inhibition Power Test

The observation results of the antagonist test of each isolate measured by observing the percentage of inhibition in vitro are presented in Table 1.

**Table 1.** Locations of various bacterial isolates isolated from the rhizosphere of the wakegi onion and the average inhibition power (%) of isolates on the growth of Foc

| Locations | Isolate Code | Inhibition power (%) |
| --- | --- | --- |
| Petobo | KP17 | 48.82 ± 3.06 |
| | KP5 | 43.28 ± 1.27 |
| Bulupontu | DB9 | 51.47 ± 1.34 |
| | DB12 | 67.66 ± 3.91 |
| | DB18 | 49.70 ± 1.36 |
| Guntarano | DG4 | 46.25 ± 2.76 |
| | DG11 | 51.19 ± 4.02 |

Inhibition power data are in mean ± standard deviation

The results of the antagonist test on the seventh day show that all local isolates of rhizosphere bacteria obtained could inhibit the growth of pathogenic mycelium from 43.28 to 67.66%. The highest inhibition power was found in DB12 isolates at 67.66% and the lowest was in KP5 isolates with an inhibition power of 43.28%. The DB12 isolate shows the highest effect, causing more than 60% inhibition of mycelium growth compared to isolates KP17, KP5, DB9, DB18, DG4, and DG11 (<60%). Thus, strain DB12 was the isolate selected to be developed because it had the best antifungal potential as indicated by a significant inhibition effect.

The inhibition test was done to obtain rhizosphere bacterial isolates that had the potential as biocontrol agents against plant pathogens. The results of the in vitro inhibition power test using the multiple culture method show that all local isolate bacteria were able to inhibit the growth of Foc pathogens, but each isolate showed different percentages of inhibition power. The difference in the inhibition effect that occurred was probably due to the different strains of the wakegi onion rhizosphere bacteria, which produced different secondary metabolite compounds, both in production (volume) and type. According to Nega [32], the production of secondary metabolites was strain-dependent, namely different strains of the same bacterial species could produce different products and types of metabolites so that the effect would be different on the target pathogen. Even though the isolates were in the same species or genus, they were not necessarily the same strain because the phenotypic character could change due to environmental effects (not static). Therefore, strain selection was an important part of the development of bacterial strains as biocontrol agents of plant diseases.

All local isolates of rhizosphere bacteria were able to form a clear zone on the PDA media (Figure 1A). Isolates of rhizosphere bacteria that grew and were able to form clear zone were indicated as isolates that had an antibiosis mechanism in suppressing the growth of pathogenic fungal colonies [33].

**Figure 1.**
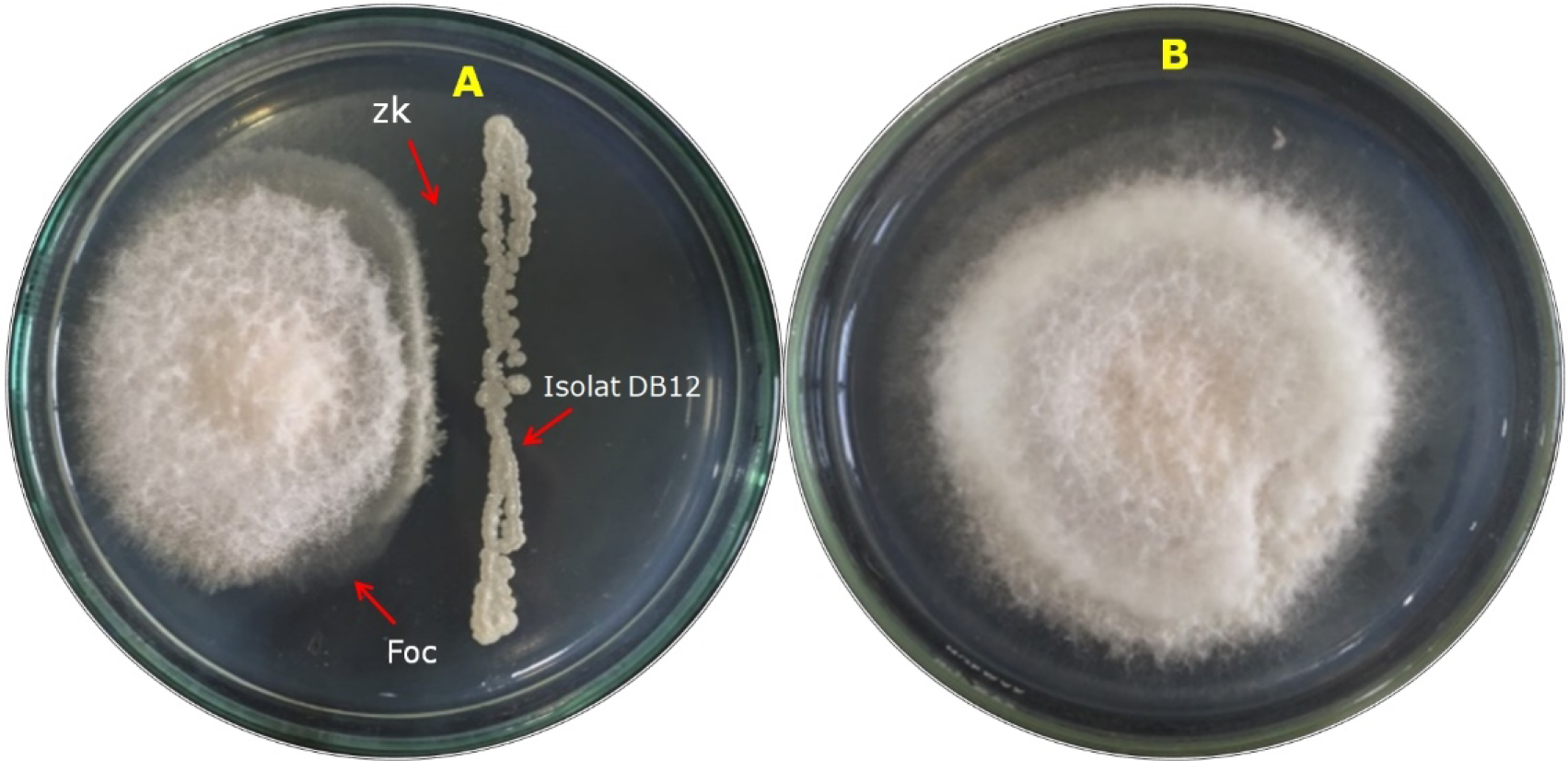
In vitro inhibition power test of DB12 isolate on the mycelium growth of *F. oxysporum*. *cepae*: clear/empty zone (zk) (A) and pathogen colony (B)

The inhibition power of local isolates on the growth of Foc mycelia was thought to lead to an antibiosis antagonist mechanism. Antibiosis was a mechanism of antagonistic bacteria by producing secondary metabolites in the form of antibiotics or antibiotic-like compounds such as lytic enzymes, volatile compounds, siderophores, and other toxic substances [34]. The mechanism of antibiosis was characterized by the formation of a clear zone between isolates of rhizosphere bacteria and pathogens on solid media (PDA). The formation of this clear zone indicated the presence of an antibiosis mechanism of the isolate against pathogens [35]. The presence of this clear zone indicated the presence of active secondary metabolites (antibiotics) secreted by isolates of rhizosphere bacteria that have spread in PDA media [13]. Secondary metabolites produced by antagonistic bacterial strains showed antifungal potential [15]. This antifungal activity was shown by the formation of a clear zone produced by the activity of antibiotic compounds [36]. The production of this antibiotic was a major step in increasing the competitiveness of rhizosphere bacteria such as *Bacillus* spp. under a limited resource environment [37]. Antibiotics were secondary metabolite products produced by an organism in small amounts that could kill or inhibit other microorganisms [36]. Toxic effects of antibiotic compounds could interfere with the normal growth process of pathogenic fungi such as changes in the shape of mycelia (malformations), as well as inhibit the growth of mycelia and spore germination [27]. The malformations in this inhibition power test were indicated by the shortening of the colonies (mycelia) that approached the rhizosphere bacteria isolates. This shortening of the colony was a form of abnormal growth in hyphae, which caused mycelia, could not develop properly on artificial media (PDA) around the rhizosphere bacterial colony.

The inhibition of the pathogenic fungi growth by rhizosphere bacterial isolates confirmed that these local isolates had the potential to produce several secondary metabolites such as antifungal compounds. Khedher et al. [38] reported that the formation of a clear zone in an in vitro inhibition power test of the mycelia growth of the *Rhizoctonia solani* fungus indicated that *B. subtilis* bacteria produced antifungal compounds. Many antifungal compounds secreted by *Bacillus* spp. have been reported, including fengisin, bacilisin, lipopeptide [39]. These secondary metabolites played an important role in the control of plant pathogens [40].

Secondary metabolite compounds produced by rhizosphere bacterial isolates to suppress the pathogenic fungi mycelia growth on PDA media were thought to be lipopeptide antibiotics compounds. Although the identity of the antibiotic was not tested in this research, the assumption was based on the ability of the lipopeptide to be heat-resistant and antifungal. In this research, the suspension of rhizosphere bacterial isolates obtained was heated before being used for the inhibition power test. Ruiz-Sanchez et al. [14] reported that lipopeptide compounds such as iturin were antifungal and heat-resistant. Malviya et al. [41] added that lipopeptides were stable to high temperature and pH, and showed high ability as biocontrol agents against various types of pathogens. Furthermore, it was added that almost all bacteria species of *Bacillus* spp. produced antimicrobial compounds were known as lipopeptides. *Bacillus* spp. bacteria were known to have a capsule-shaped structure containing a polypeptide of D-glutamic acid and are spore-producing bacteria [42]. *Bacillus subtilis* spores in the form of endospores could survive in extreme environmental conditions or were very tolerant of heat and drought [43,44]. Cyclic lipopeptides were divided into three groups, namely iturin, fengisin, and surfactin [45]. Iturin has strong antifungal activity against a wide variety of fungal and yeast pathogens and could be used as a biopesticide for plant protection. Meanwhile fengisin has antifungal activity in inhibiting the growth of various plant pathogens, especially filamentous fungi [46]. Fengisin worked by affecting the cell membrane of pathogenic fungi to change its permeability, resulting in the release of cell contents [12]. Ali et al. [15] added that the antibiotic fengisin had strong antifungal activity against filamentous fungi because it interacted with sterol and phospholipid molecules in the membrane, thereby changing the structure and permeability of fungal cell membranes.

The ability of local isolates of rhizosphere bacteria to inhibit pathogens showed that these isolates were able to act as biological control agents. A microbe was said to be a biological agent if it could inhibit the development and growth of other microbes [47]. Among isolates of rhizosphere bacteria found, isolate DB12 was the most effective bacteria strain, with the potential for inhibiting the growth of Foc pathogenic mycelium more than 60% in vitro. According to Noverizaa & Quimio [48], a microbe was said to be antagonistic if it had inhibitory activity above 60% against pathogenic mycelium. Thus, the isolate produced an inhibition zone large enough to prevent physical contact with the pathogen.

The ability to inhibit the growth of pathogenic fungi proved that DB12 isolate had the potential to be utilized and developed as a biological agent in controlling pathogens in plants. This strong inhibition power could be used as an indicator of the ability of DB12 isolates as a biocontrol agent to suppress the growth of pathogens in the field.

Furthermore, observations of colony morphology, physiological and biochemical properties of rhizosphere bacteria, as well as colony morphology and pathogenic cells were done to determine their characteristics.

### Characteristics of Local Isolates Bacteria

The macroscopic characterizations of all colonies of local isolates that have shown antibiosis activity on NA media are presented in Table 2.

**Table 2.** The morphological characteristics of isolate colonies of rhizosphere bacterial

| Characteristics | Isolate Code |  |  |  |  |  |  |
| --- | --- | --- | --- | --- | --- | --- | --- |
|  | KP17 | KP5 | DB9 | DB12 | DB18 | DG4 | DG11 |
| Shape | round | irregular | irregular | irregular | round | round | round |
| Color | dull white | beige | dull white | dull white | beige | beige | beige |
| Edge | jagged | flat | jagged | jagged | flat | flat | Flat |
| Elevation | flat | flat | flat | flat | flat | flat | Flat |
| Texture | fine | rough and dry | fine | fine | rough and dry | rough and dry | rough and dry |
| Colony size | medium | medium | medium | medium | medium | medium | medium |
| Appearance | dull | dull | dull | dull | dull | dull | Dull |
| Optical properties | opaque | opaque | opaque | opaque | opaque | opaque | opaque |
| Slimy/not slimy | not slimy | not slimy | not slimy | not slimy | not slimy | not slimy | not slimy |

The results of characteristic observations for local isolate bacterial colonies obtained from the rhizosphere of the wakegi onion were colonies in round or circular shape, dull-white or slightly yellow in color, opaque optical properties, flat elevation, and rough and dry texture with medium colony size (Figure 2A). The results of characteristics observations of local isolate colonies had similarities with the characteristics of *Bacillus* spp. bacteria described by Schaad et al. [21], Calvo & Zuniga [22], and Lu et al. [23]. In Figure 5B, it could be seen that the center of the local isolate colony contained a concentric ring-shaped circle and an irregular center border. These characteristics were similar to *Bacillus* spp. found by Calvo & Zuniga[22].

**Figure 2.**
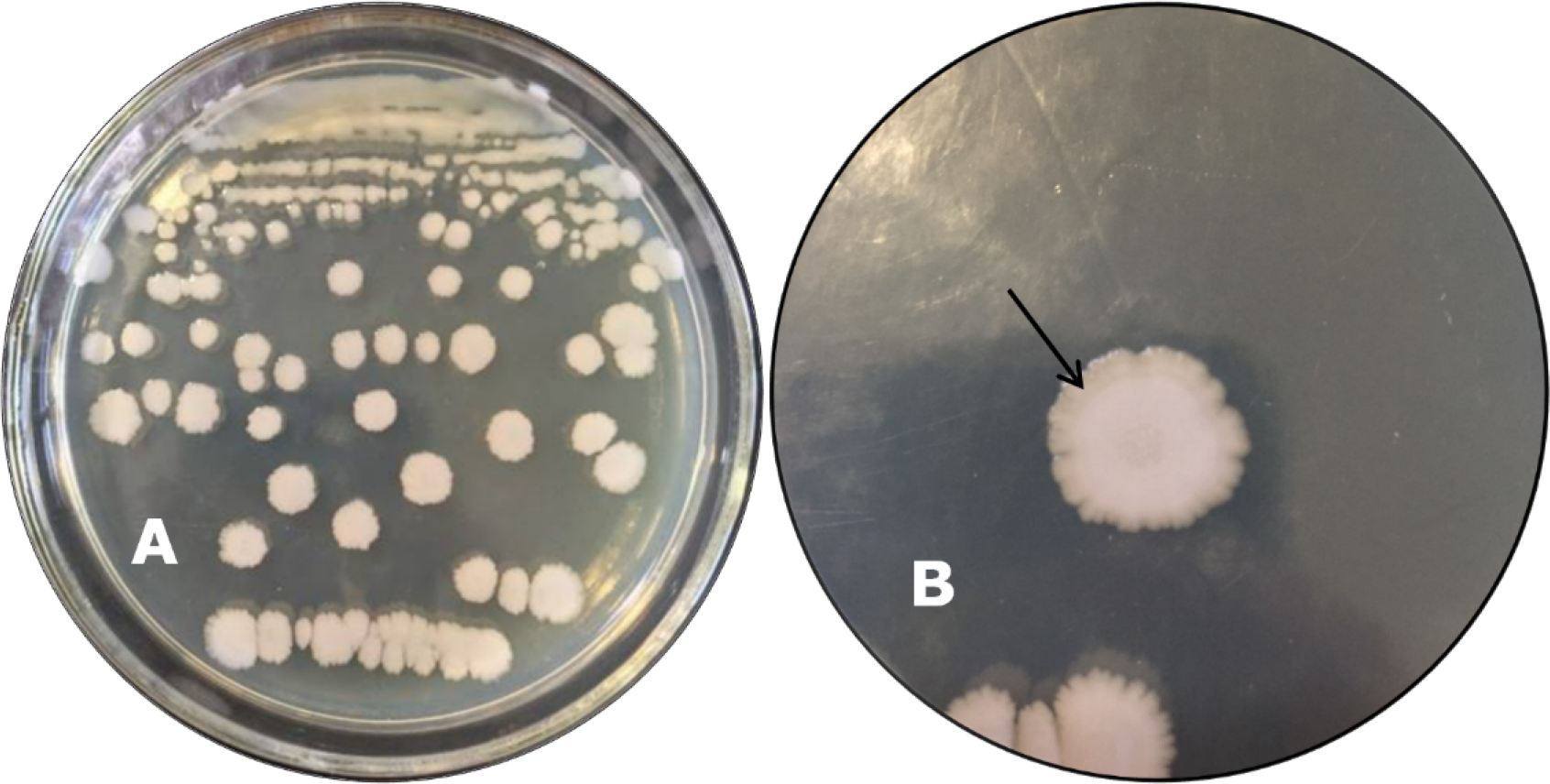
Macroscopic characteristics of bacterial isolates of DB12 (A); a single colony of local isolate (B); in the center of the bacterial colony, there is a ring-like circle (arrow)

Based on the physiological and biochemical characteristics, the local isolates were gram-negative bacteria, aerobic or fermentative anaerobic, with positive oxidation, positive catalase, did not produce fluorescent pigments, and were able to grow at a temperature of 65°C (Table 3 and Figure 3). These properties were similar to the characteristics of *Bacillus* spp. Bacteria that have been described by Schaad et al. [21], Nurcahyanti & Ayu [24], and Prihatiningsih et al. [25]. Based on the morphological characteristics of the colonies as well as physiological and biochemical properties, it was suspected that the local isolates obtained were bacteria from the *Bacillus* genus.

**Figure 3.**
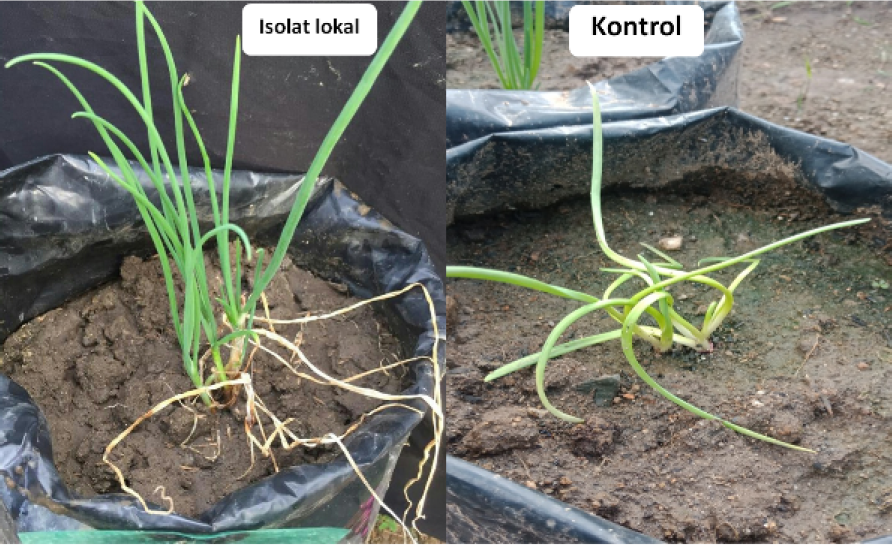
*Fusarium* wilt disease suppression test

**Table 3.**
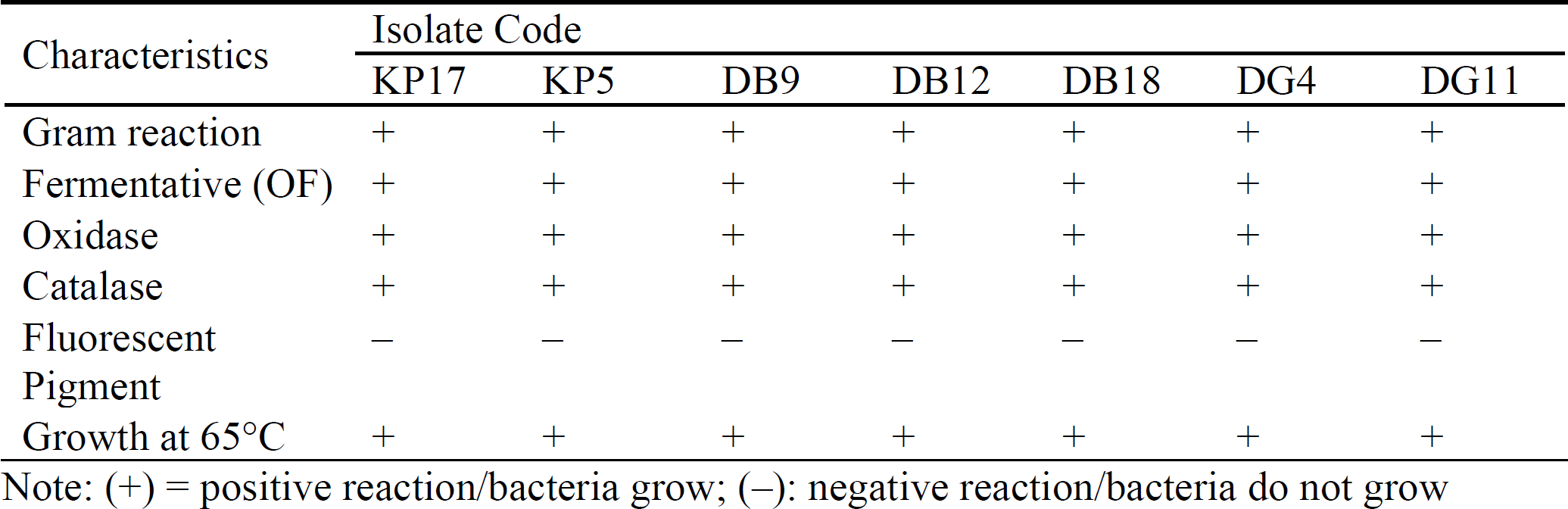
Physiological and biochemical characteristics of local isolates of wakegi onion rhizosphere bacteria

### In the Greenhouse

The incubation period of *Fusarium* wilt disease was observed daily, from the beginning of the inoculation of the pathogen until the infection occurred, which was expressed in the form of the first symptoms on the wakegi onion plant. Early symptoms appeared visibly in the form of wilt leaves that became bent or twisted starting from the top, and yellow-coloration (chlorosis). The results of data analysis show that the application of local *Bacillus* spp. isolates had a significant effect on the incubation period of *Fusarium* wilt disease (Table 4).

**Table 4.**
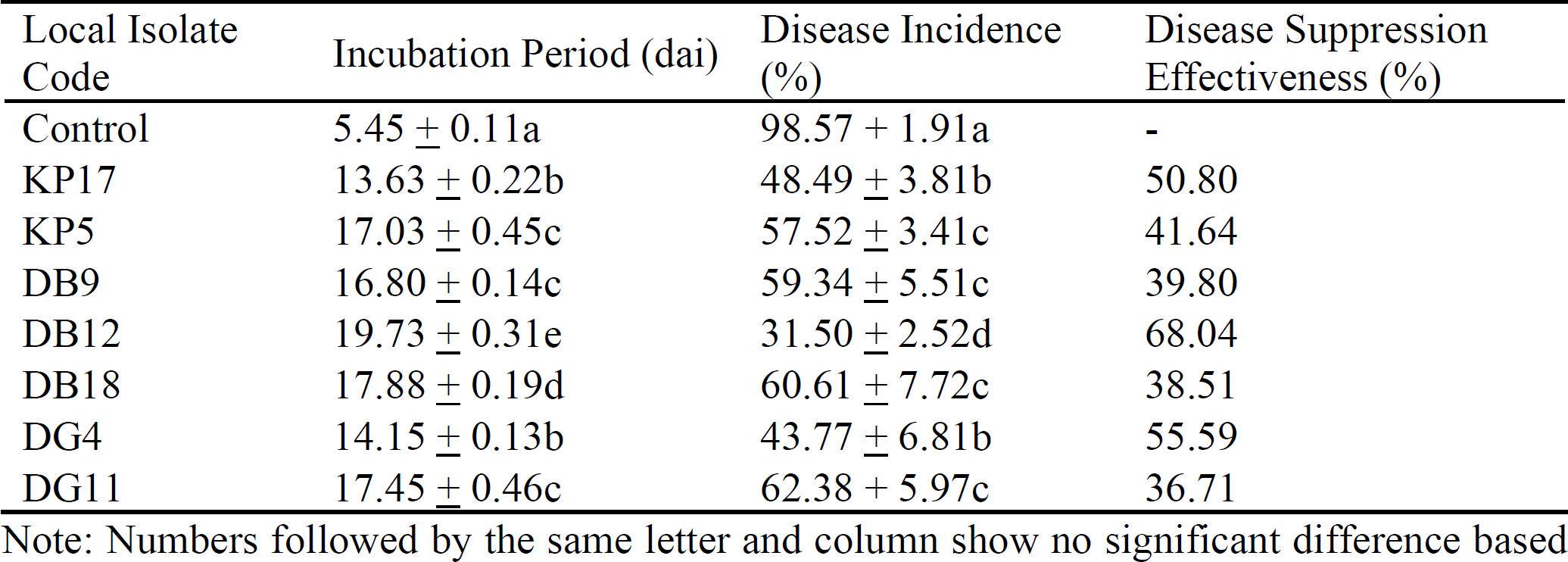
The average incubation period (dai), disease incidence (%) in various local isolates of the wakegi onion rhizosphere; incubation period data in mean ± standard deviation

In Table 4, it can be seen that the application of local isolates gave a significantly longer incubation period than the control. In the control, the incubation period required for the appearance of the first symptoms was 5.45 dai, while in the application of local isolates there was a delay in the appearance of disease symptoms with an incubation period between 8.18–14.28 dai. This shows that the application of local isolates to protect plants from pathogen attack resulted in a slower incubation period when compared to controls. The DB12 isolate had the longest incubation period (19.73 dai) and was significantly different from all tested isolates. The appearance of the first symptoms of *Fusarium* wilt disease occurred more quickly in the control due to the absence of *Bacillus* spp. bacteria as a biocontrol, such that the pathogen became more aggressive in infecting to cause disease. The absence of *Bacillus* spp. biocontrol agents caused the pathogen to infect plant roots quickly, making the incubation period shorter before symptoms appeared. The *F. oxysporum* fungus is known to be able to grow and develop in the soil and infect plant roots quickly [49].

The incubation period was longer than the control because the *Bacillus* spp. bacteria applied were local isolates obtained from the wakegi onion rhizospheres in the field so that they were more adaptable (pH, nutrition, temperature, and humidity) to the soil planted with wakegi onions in polybags. This proved that all bacteria applied were able to adapt, grow, and develop well in polybags, which was indicated by a delay in the incubation period. The ability to adapt to the environment in artificial media would accelerate microbial growth followed by an increase in the ability to produce toxins [50]; therefore, in the field it would support the ability of antagonists to suppress soil-borne pathogens. Santhanam et al. [51] stated that the use of local antagonist microbes tended to have better adaptability compared to isolates introduced from elsewhere.

The delay in the incubation period of wilt disease on wakegi onions shows that the bacteria applied were actively working in inhibiting the development of pathogens. The activity that occurred was probably a form of the mechanism of antagonistic bacteria in antibiosis in suppressing and inhibiting the growth of pathogens. According to Ali et al. [15] the antibiosis mechanism of *Bacillus* spp. played a key role in the biocontrol of plant diseases. The antibiotic activity of the *Bacillus* spp. biocontrol organisms was done by producing antibiotics that could inhibit the growth of pathogens to delay the incubation period. This was in line with Campbell’s 1. [52] statement that the antagonist mechanism of *Bacillus* spp. bacteria in an antibiosis manner tended to lead to the ability to produce antibiotics. The ability to produce antibiotics was a fundamental mechanism of *Bacillus* spp. bacteria in protecting plants from pathogen attack.

Krzyzanowska et al. [53] reported that *B. subtilis* was a bacterium that was commonly found in the soil, especially in the rhizosphere area, and it was an effective biocontrol agent against plant pathogens because of its ability to produce various kinds of antibiotics. *Bacillus subtilis* bacteria were known to produce lipopeptide compounds that were antibiotic and had a broad spectrum, such as iturin, surfactin, and fengysin [54,14,56)]. These compounds have shown potential antagonistic activity against pathogenic bacteria and fungi in both in vitro and in planta conditions to suppress disease incidence and severity [14,29].

The results of data analysis show that the application of local isolate bacteria had a significant effect on the incidence of *Fusarium* wilt disease (Table 4). In addition to delaying the incubation period, the application of local isolates was also very effective in reducing the incidence of *Fusarium* wilt disease. During observations 56 days after inoculation, the application of local isolates to suppress the incidence of disease was significantly different from the control. The average percentage of disease incidence in the application of local isolates was lower than in the control. In the control, the disease incidence reached 98.57% (the highest), while in the application of local isolates, the disease incidence that could be suppressed ranged from 38.51 to 68.04%. This proves that the application of local isolates could significantly reduce the incidence of *Fusarium* wilt in wakegi onions compared to controls. Among the tested isolates, isolate DB12 had the lowest percentage of disease incidence and was significantly different from other test isolates (Figure 3).

The success of *Baclllus* spp. local isolates in reducing the percentage of *Fusarium* wilt disease incidence was in line with their ability to delay the appearance of symptoms indicated by a longer incubation period. Every time there was a delay in the incubation period, it would generally be followed by a decrease in the percentage of disease incidence. *Bacillus* spp. bacteria especially DB12 isolate have shown the ability to act as a biocontrol agent against pathogenic fungi because it was able to delay the incubation period longer than the control, followed by the ability to suppress disease incidence to 31.50% lower than the control. This proves that *Bacillus* spp. local isolates had higher adaptability to survive and had a better chance of controlling pathogens in their place of origin than using isolates introduced from elsewhere.

The ability of *Bacillus* spp. to suppress Foc pathogens was thought to be caused by an antibiosis mechanism by producing antibiotic compounds to counter the damaging effects of these pathogens. Arseneault and Filion [57] reported that antibiosis was the most widely used mechanism of *Bacillus* spp. to control plant pathogens. *Bacillus* spp. bacteria were known to produce twelve major antibiotics including bacillomicin, mycobacillin, fungistatin, iturin, fengisin, plipastatin, and bacillin [15].

Research on the use of antibiotics produced by bacteria belonging to the genus *Bacillus* spp. as a biocontrol against pathogens has been widely reported. The results of the research by Plaza et al. [54], showed that surfactin antibiotic compounds secreted by bacterial strains belonging to the species *B. subtilis* could inhibit the mycelia growth of the fungi of *Botrytis cinerea, Sclerotinia sclerotiorum, Colletotrichum gloeosporioides*, *Phoma complanata*, and *Phoma exigua* var. *exigua*. Zalila-Kolsi et al. [58] reported that the iturin antibiotic compounds, fengisin, surfactin produced by strains of bacteria *Bacillus* spp. were able to suppress the growth of the *Fusarium graminearum* fungus. Similarly, strains of *B. subtilis* PCL1608 and PCL1612 could produce high levels of antibiotics, especially iturin A which served as the main mechanism in controlling *F. oxysporum* and *Rosellinia necatrix* [56].

In addition to using an antibiosis mechanism (antibiotic production), the biocontrol activity of *Bacillus* spp. could also be through the mechanism of parasitism (lysis of pathogenic hyphae), inhibition of enzymes or toxins produced by pathogens, competition for space and nutrients, triggering plant growth, and induction of systemic resistance [38,46]. *Bacillus* spp. isolated from soil rhizosphere has been reported to be effective in controlling various fungal pathogens [27] including *Pythium* spp., *Botrytis* spp., *Phytophthora* spp., *Fusarium* spp. [15]. Therefore, the low incidence of *Fusarium* spp. wilt disease indicates the ability of local isolates *Bacillus* spp. in producing antibiotics to suppress the growth of Foc pathogenic fungal colonies in the wakegi onion rhizosphere so that the mycelium of pathogenic fungi has difficulty growing in soil media.

## Conclusion

The isolates obtained from the wakegi onion rhizosphere could inhibit the growth of *F. oxysporum* f. sp. *cepae* colonies, a pathogen on wakegi onions, in vitro with an inhibition power between 43.28 and 67.66%. The results of the characterization of bacterial isolates from the rhizosphere of the wakegi onion lead to their identification as *Bacillus* spp. bacteria because there were similarities to *Bacillus* spp. that have been reported by previous researchers based on colony morphology as well as physiological and biochemical characteristics. The DB12 isolate of the *Bacillus* spp. bacteria had the best inhibitory ability, with the longest incubation period (19.73 dai) and the lowest disease incidence (31.50%).

